# Viral vector-driven trans-encapsidation of replicon RNAs as a rapid approach for the development of safe and economically attractive anti-enterovirus vaccines

**DOI:** 10.1101/2025.12.01.691592

**Authors:** Anna Zimina, Diana Kouiavskaia, Ekaterina G. Viktorova, Seyedehmahsa Moghimi, Konstantin Chumakov, Amy B. Rosenfeld, George A. Belov

## Abstract

Multiple enteroviruses are associated with life-threatening and economically important diseases, yet licensed vaccines are available only against poliovirus (worldwide) and enterovirus A71 (China). Both live attenuated and inactivated anti-poliovirus vaccines, while highly successful in preventing the disease, have important shortcomings. Live vaccine strains are inherently genetically unstable and can regain virulence, leading to re-emergence of paralytic disease.

Inactivated vaccine does not induce the mucosal immunity sufficient to interrupt viral transmission and is made from virulent strains, presenting a biosafety challenge. Recent alternatives, such as new vaccine strains with improved genetic stability and VLP-based vaccines, only partially address these concerns. Here, we investigated another approach to the development of anti-enterovirus vaccines based on efficient trans-encaspidation of replication-competent enterovirus RNAs coding for only the non-structural proteins (replicons) by Newcastle Disease virus vectors expressing enterovirus capsid proteins. Thus, the encapsidated replicon production is driven by effectively replicating enterovirus RNA and the NDV vector. This system is easily scalable and can be adapted to any cell culture provided it can be infected by both the enterovirus and NDV. Unlike the empty VLPs, the encapsidated replicons recapitulate the stability and antigenicity of native enterovirus particles, but cannot propagate beyond the originally infected cell. The protective efficacy of an encapsidated poliovirus replicon immunization was similar to that of the licensed Sabin vaccine strain in a murine model. This approach can easily be adapted to any enterovirus, allowing the rapid development of new, affordable vaccine candidates.

## Introduction

Enteroviruses are the most numerous group of viruses infecting humans. Currently, the genus *Enterovirus* of the family *Picornaviridae* comprises more than 300 different genotypes of human-specific viruses (1). They include well-established pathogens, such as poliovirus, Enterovirus A71, Coxsackieviruses, as well as emerging threats such as Enterovirus D68. Enterovirus infections are associated with diverse complications, including but not limited to partial or full paralysis, encephalomyelitis, myocarditis, neonatal sepsis, asthma, and could be linked to the development of type 1 diabetes (2–4).

All enteroviruses share the same genome organization and replication strategy. The viral RNA genome codes for one polyprotein and serves as an mRNA upon infection. The polyprotein undergoes co- and post-translational processing by virus-encoded proteases. The precursor of capsid proteins P1 encoded by the 5’-proximal part of the genome is cleaved *in cis* by the protease 2A from the rest of the polyprotein and is further processed *in trans* into VP1, VP0, and VP3 proteins by proteases 3C or 3CD (5, 6). The three capsid proteins assemble into pentamer intermediates, which spontaneously form empty particles consisting of 60 copies of each of the capsid proteins. These particles can either encapsidate progeny genome RNAs derived from the actively replicating pool of viral RNA or persist as VLPs (7). The final autocatalytic cleavage of VP0 into VP2 and VP4, triggered by encapsidated RNA, is the final particle maturation event required for its stability (8). Importantly, the enterovirus replication process is independent of structural proteins, so that RNAs coding for only P2-P3 proteins are fully replication-competent. Currently, the only vaccines available to control enterovirus infections are inactivated and live attenuated vaccines against poliovirus (9), and inactivated vaccines against Enterovirus A71, recently licensed in China and some other countries (10). The long history of massive administration of both oral live and inactivated anti-poliovirus vaccines provides ample data for the evaluation of challenges facing the development of any anti-enterovirus vaccine. Both vaccines confirmed their exceptional efficacy in preventing paralytic poliomyelitis, but also brought to light important problems that contribute to the difficulties facing the Global Polio Eradication Initiative. The live attenuated vaccine is relatively easy to produce and is easy to administer orally. However, as poliovirus containment measures are imposed, propagation of even attenuated polioviruses requires significant investments in biosecurity. In about one in a million recipients, live vaccine induces the disease (vaccine-associated paralytic poliomyelitis, VAPP). Upon transmission in communities, vaccine strains can lose the attenuation determinants and establish continuous circulation, giving rise to circulating vaccine-derived polioviruses (cVDPV) (11). The vaccine-derived strains now constitute the major pool of circulating polioviruses ((12), current data are available at https://polioeradication.org/circulating-vaccine-derived-poliovirus-count). The inactivated vaccine is safe, but it is costly to produce and administer, with ever more strict biosecurity measures required for propagation of wild-type infectious viruses, increasing the production cost. Another important disadvantage of the inactivated polio vaccine is its inability to induce robust mucosal immunity and prevent the spread of poliovirus in IPV-immunized populations (13, 14). To overcome the supplemental cost of polio containment, IPV composed of attenuated poliovirus strains was developed (15–17).

However, such an approach requires the development of novel manufacturing procedures and quality control reagents and addresses the biosecurity concerns only partially, since attenuated strains should also be handled under strict containment conditions.

Thus, alternative anti-poliovirus vaccines that can alleviate the safety and other limitations of the currently used vaccines are being actively explored. One approach was to further modify the attenuated strains of poliovirus to overcome their inherent genetic instability (18). However, upon their real-world deployment, environmental circulation and cases of paralytic disease associated with the improved type 2 strain have already been documented (12, 19). Another direction to increase vaccine safety is to switch from infectious virion propagation to the production of empty virus-like particles (VLPs). This can be achieved by different approaches – expression of poliovirus capsid proteins from a viral vector, which can be administered as a live vaccine (20), or production and purification of VLPs using economically viable expression systems, such as yeast, plants, or insect cells (21–23). Yet, VLPs formed by unmodified capsid proteins are structurally less stable than enterovirus virions because the final maturation cleavage of VP0 does not occur without the RNA, and they are not stabilized by the packaged RNA. VLP stability may differ between enteroviruses, and at least for poliovirus and Enterovirus A71, select amino-acid substitutions can increase VLP stability (23–26). However, stabilized polio VLPs were not immunogenic without an adjuvant in the rat potency assay (27). The loss of immunogenicity may represent a significant challenge for VLP-based enterovirus vaccine development, in addition to the requirements for optimization of VLP purification and stabilization that have to be addressed individually for each enterovirus.

Here, we developed a novel approach for the production of vaccine candidates that can be easily adapted to any enterovirus, which we name RAVE (RApid Vaccine Enterovirus) particles. Our approach produces virions encapsulating enterovirus RNAs encoding for only the proteins needed for RNA replication (replicons). Such viral particles containing replicon RNA inside retain the antigenic structure and stability of *bona fide* virions, they can initiate infection, but cannot spread beyond the originally infected cells because the RNA genome does not code for the capsid proteins. RAVE virions are safe to generate and handle since no infectious enterovirus is involved at any step of their production. In our system, the capsid proteins required for enterovirus replicon trans-encapsidation are provided upon replication of a Newcastle disease virus (NDV) vector (20). Importantly, the packaged polio replicons matched the immunogenicity and the protective efficacy of a licensed Sabin vaccine strain. This system enables rapid production of large quantities of packaged replicon particles in cell cultures approved for vaccine production, and can be used for the generation of safe vaccine candidates against any enterovirus within weeks.

## Materials and Methods

### Cells and viruses

HeLa and RD cells were grown in high-glucose modification of Dulbecco’s Modified Eagle Medium (DMEM) supplemented with 10 mM sodium pyruvate, 10% fetal bovine serum (FBS), and antibiotic-antimycotic mix (GIBCO) at 37 °C with 5% CO₂. HeLa cells were used for poliovirus propagation, replicon packaging, and plaque and immunofluorescence assays. RD cells were used for the microneutralization assay. Attenuated poliovirus strain Sabin-1 was a progeny of the US National neurovirulence reference (NA4), wild-type Mahoney (type 1) and Saukett (type 3) strains were obtained from Sanofi-Pasteur, where the strains were used for IPV manufacture. Recombinant Newcastle disease virus expressing capsid polyprotein precursor P1 and protease 3CD (NDV-polio) was generated in our lab and described in detail in (20). NDV-polio was propagated in embryonated chicken eggs as described in (28).

### Replicon encapsidation and purification

RNA coding for poliovirus type 1 Mahoney genome lacking the P1 capsid protein-encoding region (replicon) was in vitro transcribed using the MEGAscript™ T7 High Yield Transcription Kit (Invitrogen) from the linearized pXpA-P2P3 plasmid and purified as described in (29). Briefly, the RNA was extracted with phenol-chloroform, ethanol-precipitated, dissolved in nuclease-free water, and additionally purified using a Chroma Spin-100 column (Takara). This RNA was used for the first round of generation of encapsidated replicons. HeLa cells were infected with recombinant NDV-polio at an MOI of 10 and incubated at 37 °C overnight. The next day, the cells were transfected with the polio replicon RNA using TransIT-mRNA Transfection Kit (Mirus) and incubated for an additional 6 hours at 37 °C. The cells were freeze–thawed three times to release encapsidated replicon particles. The lysate was clarified by low-speed centrifugation to remove cell debris, and the supernatant containing the encapsidated replicons was collected. This supernatant was used as the input material for the subsequent propagation of encapsidated replion in place of RNA transfection.

For large quantity packaged replicon purification, four to six T-75 flasks co-infected with recombinant NDV-polio and packaged P2P3 replicon were freeze-thawed three times. Cell debris and medium were collected, resuspended in 5 ml of RSB (10 mM NaCl, 10 mM Tris-Cl, pH 7.4, 1.5 mM MgCl2), and adjusted to 1% NP-40, followed by centrifugation at 6000g for 10 min at 4 °C, and the supernatant was collected. The pellet was similarly resuspended in 3 ml of 1% NP-40 in RSB and clarified by centrifugation three times. The combined supernatants were clarified by an additional centrifugation at 17000g for 10 min at 4 °C. Upon addition of sodium dodecyl sulfate (SDS) to a final concentration of 1%, the supernatants were centrifuged at 40,000 rpm for 4 hours in a Beckman SW41 Ti rotor. The pellet was resuspended in 0.5 ml TE buffer and subjected to a second round of ultracentrifugation under identical conditions to remove residual SDS. The final pellet was resuspended in 1 mL TE buffer, passed through a 0.22 µm filter, and stored at −80 °C. The same purification procedure was used for the preparation of Sabin-1 stock for immunization.

### Poliovirus and encapsidated replicon titration

Since encapsidated replicon particles cannot generate plaques, for consistency, titers of both Sabin-1 poliovirus and encapsidated replicons were determined in an immunofluorescence assay as fluorescence-forming units (FFU)/ml.

HeLa cells seeded on cover slips were infected with serial dilutions of encapsidated replicon or Sabin-1 virus and incubated at 37 °C for 4 h or 34 °C for 6 hours, respectively (so that cells are analyzed during the first replication cycle, before the virus spread can occur). Cells were fixed with 4% paraformaldehyde in PBS and permeabilized with 0.2% Triton X-100 in PBS for 5 min at room temperature and washed three times with PBS. Cells were blocked in 3% low-fat dry milk in PBS for 30 min, followed by incubation with anti-2B mouse monoclonal antibody dilution in a blocking buffer for one hour at room temperature. After three PBS washes, cells were incubated with Alexa-555 Fluor-conjugated anti-mouse secondary antibody (Invitrogen) diluted in a blocking buffer and Hoechst 33342 nuclear stain for 1 hour. Following three final PBS washes, coverslips were mounted using Fluoromount-G medium (Invitrogen). Fluorescence-based titers were determined by quantifying fluorescent-positive cells at 40x magnification. The FFU/ml titers were calculated based on the field-of-view area (0.26 mm²) and the number of positive cells averaged from 10 individual images per sample.

Poliovirus type 1 Mahoney and type 3 Saukett titers were determined in a standard plaque assay. HeLa cells were seeded in a 6-well plate to form an 80-90% confluent monolayer. Serial 10-fold virus dilutions were prepared in DMEM, and 200 µL of each dilution was added per well. Mock infections were performed with 200 µL DMEM alone. Plates were incubated for 30 min at room temperature on a rocking platform for virus adsorption, after which the inoculum was aspirated, and cells were overlaid with 4 mL of 0.5% agarose in DMEM supplemented with 2% FBS and antibiotic-antimycotic solution. The plates were inverted and incubated at 37 °C for 48 h, after which the agarose layer was removed, and the cells were stained with crystal violet. The same plaque assay procedure was used to assess the preparations of packaged replicons for the emergence of recombinant viruses.

### Animal immunization and challenge protocols

All animal experiments were performed according to protocols approved by the White Oak consolidated animal program at the FDA. TgPVR21 transgenic mice expressing human poliovirus receptor CD155 were obtained from the Central Institute for Experimental Animals (Tokyo, Japan). Five-week-old mice were immunized intramuscularly with the indicated doses of packaged replicons or Sabin-1 vaccine strain resuspended in 50µl of PBS without any adjuvants. Blood samples were collected from the submandibular vein pre-immunization and 3 weeks after the first immunization. For the vaccine protective efficacy test, mice were challenged one week after the boost dose intramuscularly into the right leg with wild-type poliovirus type 1 Mahoney or type 3 Saukett strain at a dose equivalent to 20 PD_50_ (Paralysis Dose 50, about 2E5 and 2E7 PFUs for poliovirus type 1 Mahoney and type 3 Saukett, respectively). Mice were observed for 14 days and were humanely euthanized when signs of paralysis became evident.

### Neutralizing antibody analysis

Anti-poliovirus neutralizing antibody in sera of immunized animals was determined by a standard microneutralization assay with RD cells in 96-well plates, titers were calculated by the Karber formula according to (30, 31).

### Statistical analysis

The GraphPad Prism software package was used for statistical analysis and graphical representation of data. Pairwise comparison of datasets was performed with the unpaired t-test with Welch’s correction, variances were compared using the F test.

## Results

### The enterovirus replicon encapsidation system based on capsid proteins provided by an NDV vector is safe and efficient

We previously developed an NDV-vectored vaccine for mucosal administration and production of poliovirus VLPs directly in the cells of a vaccine recipient (20), and recently evaluated a similar approach for the vaccine against enterovirus D68 (manuscript in preparation). These vectors encode enterovirus capsid protein precursor P1 and protease 3CD required for its processing as separate genes in the NDV genome (Fig. 1A).

**Figure 1.**
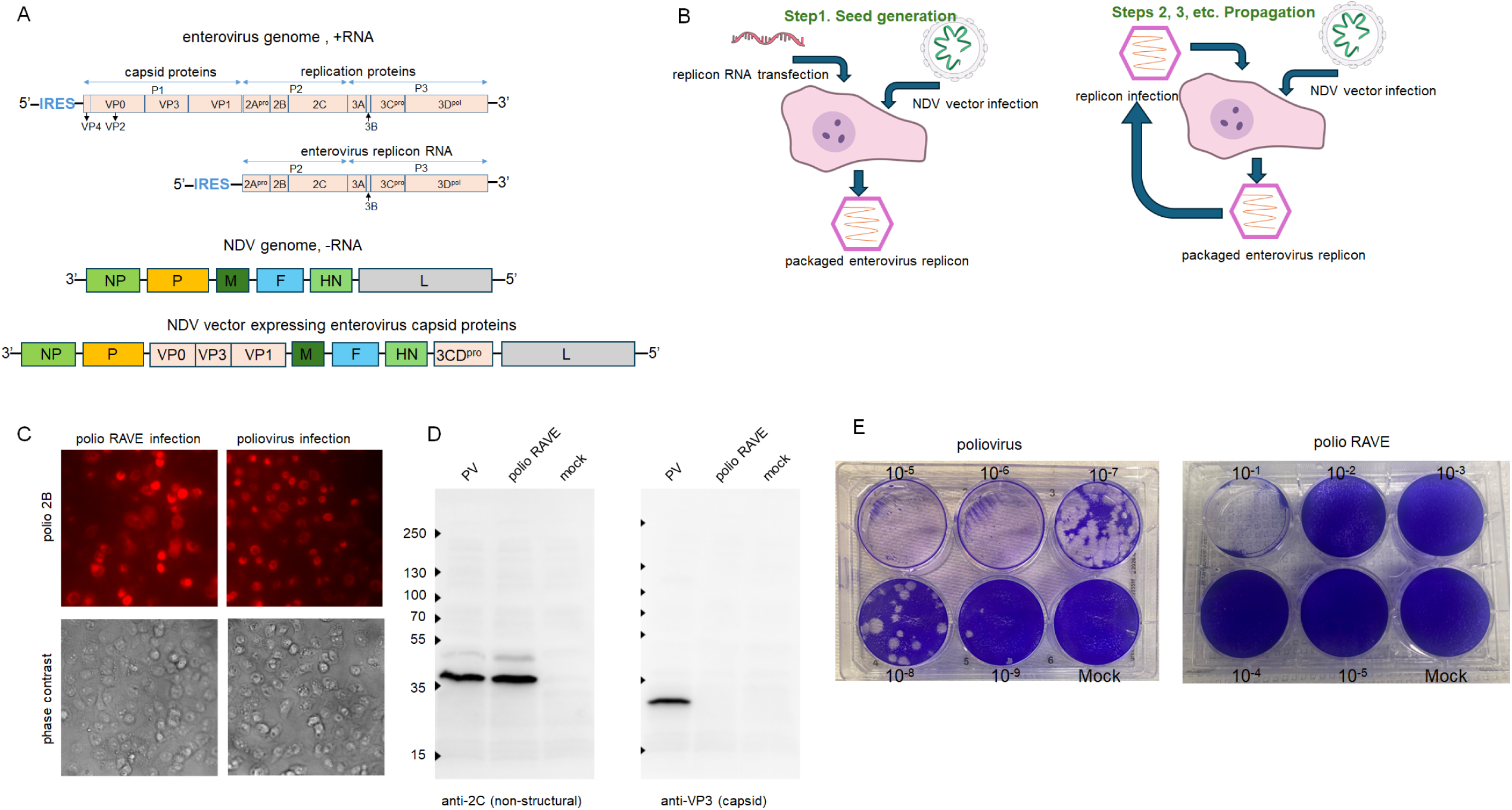
Effective propagation of encapsidated enterovirus replicons. **A.** Schemes of enterovirus genome and replicon RNA lacking the capsid protein coding sequence (top), and NDV and NDV-based vector for expression of enterovirus VLPs (bottom). **B.** Propagation of encapsidated enterovirus replicons. The figure is prepared using NIH Bioart templates. **C.** Encapsidated enterovirus replicons are infectious. HeLa cells were infected with an MOI of 10 of either purified polio RAVE or poliovirus type 1 Mahoney viruses and assessed for the expression of non-structural protein 2B at 4 h p.i. **D.** The replication efficacy is the same for the encapsidated replicon and poliovirus. HeLa cells were infected (or mock-infected) with an MOI of 50 of poliovirus type 1 Mahoney or polio RAVE infectious particles, and incubated for 4 h p.i. Cell lysates were analyzed in a Western blot assay with anti-2C (replication protein, left panel) and anti-VP3 (structural protein, right panel) antibodies. **E.** Encapsidated replicons are propagation-deficient. Plaque assays with polio RAVE virions or poliovirus type 1 Mahoney. Dilutions of the original stocks are shown. Note the destruction of the cellular monolayer only at the lowest dilution of the RAVE preparation, and the absence of plaques typical for control enterovirus titrations.

It has been directly shown for many enteroviruses that RNAs lacking capsid protein precursor coding sequence are replication competent, with a caveat that while in most members of the *Enterovirus* genus, the *cis*-acting replication element (*cre*) is located in the coding region of a non-structural protein 2C, in some of them this genetic element is located in the sequence coding for capsid proteins (32–34). In the latter case, duplication of *cre* outside of the structural protein region is required to generate a viable replicon (35).

We previously described trans-encapsidation of poliovirus replicons coding for a luciferase gene in place of capsid proteins by transfecting the replicon RNA into cells infected with the NDV vector coding for capsid proteins of poliovirus type 1 Mahoney (29).

Here, we evaluated this system as an alternative for the development of anti-enterovirus vaccines. Schematically, in the first round of replicon packaging (seed generation), cells infected with an NDV vector coding for the capsid proteins of the enterovirus of interest are transfected with an enterovirus replicon RNA coding for the replication proteins P2P2 only, generating the first yield of packaged replicons (Fig. 1B, left). This seed serves to start subsequent amplification cycles where the cells infected with the NDV vector are superinfected with the packaged enterovirus replicons, a process much more efficient and inexpensive than RNA transfection (Fig. 1B, right). The resulting packaged replicons (RAVE virions) can be purified and handled similarly to stable *bona fide* enterovirus virions. We generated purified encapsidated polio RAVE preparations with the titers of ∼E8 of infectious units per ml, which is similar to the yield of routine poliovirus propagation. These packaged replicons efficiently infected cells and initiated one round of infection (Fig. 1C). The efficacy of infection was the same as for virions with the full genome RNA, as evidenced by the similar production of the non-structural proteins in cells infected with the same MOI of polio RAVE and poliovirus type 1 Mahoney, and, as expected, no structural proteins were detected upon infection with RAVE virions (Fig. 1D).

Enteroviruses are notoriously recombination-prone, and both replication-dependent and replication-independent recombination mechanisms have been described (36–38). Thus, a recombination between enterovirus replicon RNA and the RNA coding for the capsid proteins expressed from the NDV vector is conceivable, even though by design they don’t have homology regions that may promote recombination. We thoroughly investigated all of our preparations of encapsidated replicons in a plaque assay that detects propagation-competent enteroviruses, and never observed such recombinants, even though they should have a selective advantage if they emerge. The plaque assays with polio RAVE virions showed the disruption of the cellular monolayer at lower dilutions, as expected from the high MOI at high virion concentrations, but never generated spreading plaques indicative of a recombinant virus (Fig. 1E).

Thus, encapsidated enterovirus RNAs coding for only non-structural proteins are fully infectious, indicating that they likely retain the antigenic properties of native enterovirus virions, but cannot spread beyond the first infected cell.

### The immunogenicity of the packaged polio replicon is similar to that of the vaccine Sabin-1 strain

We immunized groups of six transgenic TgPVR21 mice expressing the human poliovirus receptor (39, 40) with 1E5 FFU of poliovirus type 1 Sabin-1 vaccine strain (positive control), and with either 1E5, 1E6, or 1E7 FFU of polio RAVE intramuscularly, according to a standard procedure of poliovirus vaccine evaluation in a murine model (41, 42). A negative control group was sham immunized with PBS (Fig. 2A). Three weeks after the immunization, neutralizing antibodies were detected in two animals immunized with the Sabin-1 vaccine strain, and in five animals immunized with polio RAVE at the 1E7 dose, but not at lower doses, and not in the PBS control group. Interestingly, while the geometric means of neutralizing antibody titers from the animals immunized with Sabin-1 and polio RAVE virions were not statistically different, in the former case, the titers varied significantly, while in the latter, they showed a much more uniform distribution (Fig. 2B). We also assessed the antibody response against viral proteins in a Western blot assay. Lysates from HeLa cells infected with poliovirus type 1 Mahoney (or mock-infected control) were probed with the sera of immunized animals as a source of primary antibodies. Animals immunized with either Sabin1 or 1E7 polio RAVE virions showed a similar strong response against the protein of the molecular weight slightly less than 35 KDa, which corresponds to the structural protein VP1 (34 KDa), the major poliovirus immunogen (Fig. 2C).

**Figure 2.**
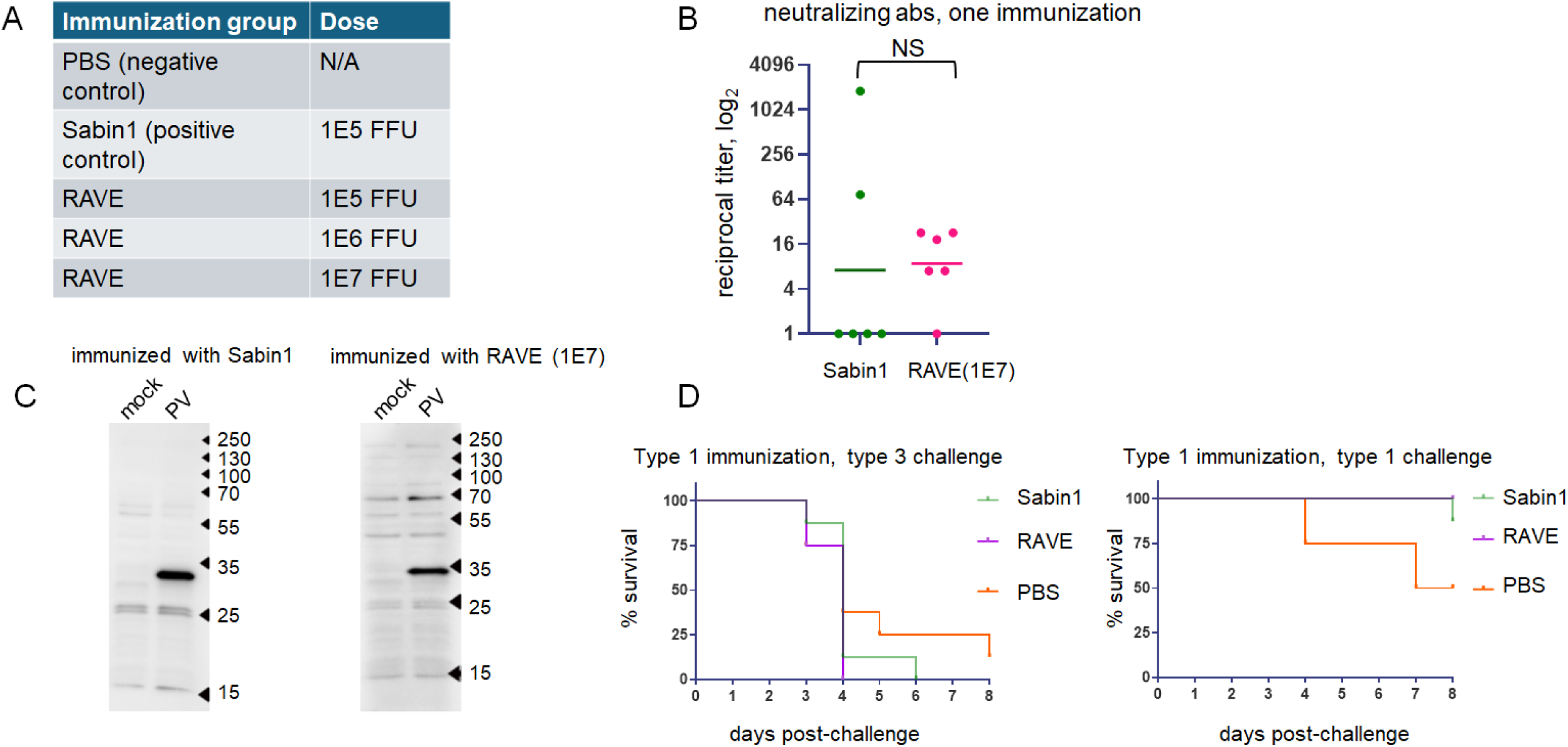
Encapsidated enterovirus virions induce a protective immune response. **A.** Immunization groups for the analysis of immune response against polio RAVE virions**. B.** Analysis of sera of animals after one immunization for anti-poliovirus neutralizing antibodies in a microneutralization assay. Data are shown for individual animals as reciprocal titers, horizontal lines indicate the geometric mean. Variances of data sets are significantly different (F(5,5)=6306, p<0.0001) **C.** Representative western blots showing similar antibody response upon immunization with Sabin1 poliovirus vaccine strain and polio RAVE virions. Lysates for Western blot were prepared from HeLa cells infected (or mock-infected) with poliovirus type 1 Mahoney. **D.** Survival of mice immunized with either control Sabin1 vaccine strain or polio RAVE virions with type 1 Mahoney capsid proteins challenged with poliovirus type 3 Sauket (heterotypic challenge, left panel), or poliovirus type 1 Mahoney (homotypic challenge, right panel).

To evaluate polio RAVE protective efficacy, transgenic TgPVR21 mice were immunized with 1E7 polio RAVE or 1E5 of Sabin-1 vaccine strain (positive control) infectious particles, or sham-immunized with PBS (negative control) according to a prime-boost scheme with a three-week interval. One week after boost immunization, the animals were challenged with poliovirus type 1 Mahoney (homotypic challenge) or poliovirus type 3 Saukett (heterotypic challenge).

Neither Sabin-1 nor polio RAVE Immunization was protective against type 3 challenge. One animal succumbed to poliovirus type 1 Mahoney challenge in the control Sabin1-vaccinated group, and all polio RAVE-immunized animals were fully protected (Fig. 2D).

## Discussion

The development of vaccines represents a classic “fast, good, and cheap” triangle problem where one has to be content with only two of the three desired qualities at a time. The traditional trial-and-error approach gave us effective and affordable vaccines, but they often took decades to develop. Recent progress with mRNA vaccination paved the way for rapid development of effective, but costly vaccines, and fast and cheap is often the domain of veterinary vaccines, where rapid livestock turnover is permissive for suboptimal vaccines. We evaluated a novel approach that can attain the sweet spot of fast generation of economically attractive and effective vaccines, which can be applied to any enterovirus, and with minor modifications to many other positive-strand RNA viruses.

The essential idea is to generate virions structurally (and therefore antigenically) indistinguishable from *bona fide* enterovirus particles, which, however, are propagation-defective. Previously, similar encapsidated poliovirus replicon particles were shown to stimulate short-term broad protection against respiratory infections upon intranasal administration in a murine model, which was attributed to the upregulation of innate anti-viral responses (43). Here, we demonstrate a new highly efficient system for the production of encapsidated replicons, suitable for industrial production, and evaluate their specific immunogenicity and protective efficacy as a vaccine. Our system consists of an enterovirus replicon RNA coding for only non-structural proteins, and an NDV vector supplying the enterovirus capsid proteins *in trans*. Both components are easy to generate for any enterovirus using standard cloning techniques. In our experiments, we used a previously developed NDV vector that expresses both the poliovirus capsid protein precursor P1 and 3CD protease. However, in principle, for the enterovirus replicon packaging, NDV vectors expressing only the P1 insert should be sufficient, as the protease can be provided by the replicon itself. This may further simplify the development of novel vaccines. This system does not involve propagation of a fully infectious pathogenic enterovirus at any stage. Moreover, since it relies on an efficiently replicating enterovirus replicon and NDV vector, it is similar to a regular virus propagation and can be easily scaled up. The NDV strain LaSota we used as a vector is a non-pathogenic strain used worldwide as a live vaccine for poultry vaccination, and its production and handling do not require strict biosecurity measures (44). Moreover, in our system, it is present only at the production stage and is inactivated by the detergent treatment and eliminated from the final product, which consists only of replicon RNA packaged in enterovirus capsids. The capacity of NDV to infect diverse types of avian and mammalian cells makes this system easily adaptable to cell cultures approved for vaccine production, provided that these cell lines are also susceptible to infection with the enterovirus particles. In this regard, the Vero cell line, broadly used for vaccine production (45), is suitable for the development of RAVE vaccines against diverse enteroviruses.

Important safety questions exist with all replication-competent vaccines. The fully infectious virus, if it emerged via recombination of the enterovirus replicon and the RNA coding for capsid proteins, would immediately outcompete the replicons since it can independently spread between cells. Yet, even though enteroviruses are highly recombination-prone (36–38, 46, 47), we never detected such recombinants. This is likely explained by the absence of sequence homology and the spatial separation within infected cells of the replicating enterovirus RNA pool from mRNAs expressed from an NDV vector. The complexity of the process is another barrier to such recombination, where the capsid protein sequence should be precisely inserted at a correct distance from the 3’ end of the enterovirus IRES and in-frame with the sequence coding for non-structural proteins. As a further safeguard against recombination, it may be possible to express individual capsid proteins from individual RNAs, as it has been shown that no uncleaved precursor is required for their assembly into VLPs (48). In any case, testing propagating pools of packaged replicons for a fully infectious virus is straightforward and can be easily implemented during industrial manufacturing. Another concern is a possible recombination of a replicon with an enterovirus currently circulating in the environment. This seems realistic only upon oral or intranasal administration of RAVE vaccines. Even if the recombination occurs, it might recreate the virus only with the capsid proteins already in circulation, i.e., it should not generate any new threats. The possibility that the replication proteins derived from the replicon would confer increased pathogenicity to the recombinant virus, while cannot be formally dismissed, seems remote. The intramuscular administration of packaged replicons, as a standard inactivated vaccine, should minimize the recombination concerns. One can also envision the adoption of packaged replicons for the production of inactivated vaccines, using the protocols established for fully infectious virions, which should at least in part alleviate production costs associated with biocontainment.

Upon intramuscular administration, the polio RAVE virions could fully protect transgenic mice from homotypic but not heterotypic challenge, similar to an approved poliovirus vaccine strain. In our assays, the immune response elicited by the packaged replicon was virtually indistinguishable from that induced by the intramuscular administration of the Sabin1 strain. It should be noted, though, that for the induction of a similar level of immune response, a much higher dose of polio RAVE virions was required than that of the fully propagation-competent live vaccine strain.

Our preparation of packaged replicons likely contained empty VLPs generated by the NDV vector. In the case of poliovirus, such VLPs were unlikely to negatively impact the immune response, as poliovirus VLPs are relatively stable and were shown to elicit the protective response even without additional stabilization measures (49). But it is possible that in the case of other enteroviruses, VLPs, which are not stabilized by the packaged RNA, may transition into non-protective antigenic conformation, resulting in suboptimal immune response. Accumulation of VLPs is inevitable upon any enterovirus propagation (50, 51), and the purification of complete virions is a standard procedure in inactivated poliovirus vaccine production (52). These established techniques can easily be applied to the purification of packaged replicon particles. In this communication, we’ve explored the intramuscular delivery of the encapsidated replicons, similar to the standard immunization with the inactivated anti-poliovirus vaccine. However, their virus-like stability allows for their oral or respiratory mucosal application, as a stand-alone regimen, or in addition to the intramuscular delivery, opening the possibility of the induction of a more comprehensive immunity, which, in the context of polio vaccination, in addition to protection against paralysis, could also at least temporarily block virus transmission. However, whether such mucosal immunization can induce durable virus-specific protection, as opposed to the previously reported short-term stimulation of innate immunity (43), requires additional research to achieve a stable high delivery rate of the propagation-defective virions to the susceptible cells in the mucosa.

In summary, we developed a highly scalable, economically attractive system for the production of packaged enterovirus replicons, which recapitulate the stability and antigenic properties of *bona fide* enteroviruses, but are safe to administer as vaccines without additional precautions. This system enables the development of vaccine candidates, suitable for industrial production, within weeks against any existing or emerging enterovirus threat.

## Acknowledgments.

The work was supported by the NIH grant R21AI153976 to GAB. AZ was supported by the Basil & Anne Hatziolos Scholarship Fund for Veterinary Medical Research.

